# Ifenprodil Intracerebrally Offers Neuroprotection against 6-OHDA-Induced Toxicity in SD-Rats via Enhancing Autophagy Function

**DOI:** 10.1101/2021.07.28.454206

**Authors:** Xinyu Zhao, Fugang Tian, Chunmin Guo, Xin Yu

**Affiliations:** School of Pharmacy, Key Laboratory of Molecular Pharmacology and Drug Evaluation (Yantai university), Ministry of Education, Collaborative Innovation Center of Advanced Drug Delivery System and Biotech Drugs in universities of Shandong, Yantai university, Yantai, China; School of pharmacy, Yantai university, Yantai, 264005, P.R. China.

**Author notes:** Corresponding author: Xin Yu. School of pharmacy, Yantai university, Yantai, 264005, P.R. China.

**Keywords:** NMDA receptor, glutamate, autophagy, Parkinson disease, Ifenprodil, Intracerebral

## Abstract

The progressive decline of dopamine neurons in the substantia nigra is the main pathogenic change in Parkinson’s disease (PD). Studies have found that excessive excitement of glutamatergic neurons causes intracellular calcium overload and induces autophagy impairment, which is one of the main mechanisms of dopamine neuron damage. The neuroprotective effect of Ifenprodil against 6-OHDA-injured mice was studied in this study. Ifenprodil was administered intraperitoneally (i.p.) or intracerebrally to rats who had a nigral-striatum pathway lesioned by 6-OHDA stereotactic brain injection. The ability to move was evaluated. The survival of dopamine neurons in the nigral was determined using HE staining, while TH-positive expression was measured using immunohistochemistry. Western Blot was used to examine the expression of CaM protein and light chain 3 (LC3), Beclin-1, BNIP3LNix, and p62. The results revealed that Ifenprodil improves motor function in 6-OHDA rats, and intracerebral injection is more effective than systemic administration. The same results also found in HE and IHC. Ifenprodil enhanced LC3II, BNIP3LNix, and Beclin-1 while decreasing p62, p-CaMKII, and β-Ca expression. In addition, Ifenprodil reduced the activation of microglia caused by 6-OHDA. Overall, the findings imply that Ifenprodil intracerebrally may protect against Parkinson’s disease via modulating autophagy-related proteins during 6-OHDA-induced toxicity.

## 1 Introduction

Parkinson’s disease (PD) is a symptomatic progressive neurodegenerative disorder characterized by motor and non-motor symptom complexes, as well as pathologically defined selective death of dopaminergic neurons in the substantia nigra(Davie, 2008; Lang and Lozano, 1998; Takeda et al., 2021). Pathological alterations in glutamatergic systems are also observed throughout PD, in addition to the dopaminergic nigrostriatal pathway. The N-methyl-D-aspartic acid (NMDA) receptor is a glutamate receptor whose malfunction is linked to the pathogenesis of PD(Su et al., 2019). It is characterized by high Ca^2+^permeability, neuronal transmission at most excitatory synapses, and overstimulation in the nigra-striatum pathway, which results in nigra neuronal damage and PD(Cali et al., 2014). The NMDA receptor is a heterotetrametric structure made up of two NMDA receptor 1-and two NMDA receptor 2-subunits(Mori and Mishina, 1995). Its signaling is complex, and the consequences might vary depending on the subunit composition of the NMDA receptor(Ahmed et al., 2011). According to reports, the NMDA receptor subtype 2B (NR2B) is involved in excitotoxicity in PD(Zhu et al., 2016), resulting in cell death and calcium overload. Although the activation of NR2B has been implicated in PD, its specific physiological function remains unknown(Hopfner et al., 2019; Masilamoni and Smith, 2018; Wang et al., 2020; Zhang et al., 2019).

Autophagy is a dynamic cellular mechanism in the turnover of proteins, protein complexes, and organelles(Anglade et al., 1997; Lynch-Day et al., 2012b; Yang and Mao,2010). Autophagy is required for brain homeostasis, and its impairment has been related to PD(Lu et al., 2020; Lynch-Day et al., 2012a). It is believed that calcium overload may play a role in autophagy impairment, which is regarded as the important pathway of the mechanisms leading to neuronal death. The role of autophagy in PD has been hypothesized as NMDA receptor activation causes a delayed inhibition in autophagic flux(Plowey and Chu, 2011). This hypothesis, however, remains elusive in PD. It is well established that glutamate receptor stimulation affects autophagy activity both in vivo and in vitro. However, it is debatable whether increased autophagy is a protective response or whether defective autophagy contributes to excitotoxic damage(Moors et al., 2017). Several investigations on PD have found that decreased LC3-I to LC3-II transition is accompanied by increased SQSTM1/p62, implying an autophagic flux inhibition, and deficient autophagy has been linked to excitotoxic mortality in these situations(Itakura and Mizushima, 2011; Law and Kim, 2014; Ney, 2015; Shvets et al., 2008). Other research suggests that excitotoxic damage is caused by activated autophagic degradation rather than suppressed autophagy.

According to various immunostaining of cultures from different brain areas, NMDA receptor antagonists can be utilized in the treatment of PD to minimize excitotoxicity. We suggest 6-OHDA-injured rats as our animal model (Lai et al., 2003; Meshul et al., 2002), to study on activated-excitotoxic pathway NMDA receptor-mediated leading vulnerability on dopaminergic neurons in the midbrain and the striatum. However, it is yet unknown how NMDA receptor antagonists work in the brain or whether they can slow the progression of PD.

The link between NMDA receptor activation and autophagy in 6-OHDA-injured rats was studied. We explored the role of NR2B in the modulatory effect of basal NMDA receptor activity on the function of autophagy using Ifenprodil, a selective NMDA receptor antagonist. Additionally, we further investigated the influence of Ifenprodil action in different brain regions.

## 2 Materials and methods

### 2.1 Chemicals

Antibodies for microtubule-associated protein 1 light chain 3B (LC3B), SQSTM1/p62, Beclin-1, bax, bcl-2, BNIP3L/Nix, PARP, and Tyrosine hydroxylase (TH) were purchased from Cell Signaling Technology (Beverly, MA, USA). Antibodies for NR2B, NR2B (phospho S1303, phospho T1472), NMDAR1, NNMDAR1 (phospho S890), ibal-2, psd95, CaMKII, phosphor, CaMKII were obtained from Abcam (Cambridge, UK). α-Ca, β-Ca were acquired from Thermo Fisher Scientific (Carlsbad, CA).

### 2.2 Drugs

Ifenprodil was received from Med CehemExpress (MCE, Qingdao, China), and was dissolved in sterile saline followed by filtration via sterile filters (Millipore Millex) for animal injections (2.56mg/kg, i.p. and 10 nM for brain injection), and dissolved in 0.9% NaCl for systemic administrations. For in vitro investigations, the compounds were solubilized in dimethyl sulfoxide (DMSO) to a final concentration of 10 mM and freshly diluted in the medium before use.

### 2.3 Animals

Animal experiments were conducted using 6-week-old Sprague Dawley rats (SPF) purchased from Pengyue Jinan. Rats were habituated to the animal facility for one week in animal housing that was kept at 23 °C with 12:12-hour light-dark cycles. Ethical approval for this study was obtained from the Care and Use of Experimental Animal Research Committee in Yantai University (APPROVAL NUMBER: YT-YX-2019-0097).

### 2.4 Medial forebrain bundle 6-OHDA lesioning

The 6-OHDA lesioned Parkinsonian rat model was performed as described in previous works(Pallini et al., 1997). In short, unilateral 6-OHDA lesion of the nigrostriatal pathway was performed with the rats under isoflurane anesthesia by stereotaxic injection of 6-OHDA HBr [(4 μg in 5 μl) per rat; dissolved in ascorbate (0.02% in NC as a vehicle)] over a period of 5 min at coordinates: AP −4.0; ML −1.65; and DV −8.0 from Bregma. The 6-OHDA-lesioned rats were given apomorphine (APO) (0.5 mg/kg) three weeks following injection. If the APO-induced rotation rate of 6-OHDA lesioned rats was greater than 7 revolutions per minute, they were selected.

### 2.5 Microinjection

Microinjection was performed as described previously(Hagg and Varon, 1993; Ragozzino et al., 2002). A rat brain atlas was employed as a reference, and stereotaxic coordinates were determined using the bregma. In brief, inserted different specifications cannula into different locations of the brain and ensure the cannulas were anchored to the skull using screws and dental cement. The guide cannula (4.0 mm) was inserted at the striatum (stereotaxic coordinates: anteroposterior, −0.26 mm to bregma; lateral, ±4.0 mm), and an injection cannula was inserted into the guide cannula (4.5 mm below the surface of the skull). The guide cannula (8.0 mm) was inserted at the midbrain (stereotaxic coordinates: anteroposterior, −4.5 mm to bregma; lateral, 1.65 mm), and an injection cannula was inserted into the guide cannula (8.5 mm below the surface of the skull). After the cannula fixing, In the guide cannula, a dummy cannula was placed. During injection days, cannulas were removed and injection cannulas attached to 10 μl Hamilton were inserted into guide cannulas. Following drug infusion (velocity= 0.25μl/min, 2μl), injection cannulas were left in place for 8 min.

The obtained concentration on the cell was used in brain treatment. Ifenprodil were delivered by cannula injection every day which persisted for at least 28 days. The cannula placements were histologically validated after behavioral assessment. Based on the region designated by the brain atlas, the successful positioning of the probe in the striatum was determined.

### 2.6 Behavior test

The following behavior tests were performed after every single administration in a half hour.

#### 2.6.1 Open field test

As previously reported(Gould et al., 2009), rats were placed in the open field chamber [50 × 50 × 70 cm (W × D × H)] and permitted to wander freely for 10 min. After each experiment, the open field was cleaned, and all tests were conducted in lower light. To quantify locomotor activity, tests were videotaped and total movement distance was calculated using a video tracking software system (SMART 3.0: Panlab, U.S.A.) to assess locomotor activity. All pencil field test was performed between 08.00 am–12.00 am.

#### 2.6.2 Rotarod test

The rotarod unit consisted of a revolving spindle (7.3 cm diameter) and four independent compartments that could test four rats at the same time, as previously reported(Hamm et al., 1994; Shiotsuki et al., 2010). After three successive days of once-daily training (increase the speed at a uniform speed within 5min, the maximum speed is 15rpm), recorded data on the last day in a test session. The duration each rat spent on the revolving bar was measured across three trials at 30 min intervals, with a maximum test time of 180 s. Over the three test trials per rat, data were shown as the meantime on the revolving bar.

### 2.7 Perfusion fixation

After the behavioral tests were completed, rats were sedated with pentobarbital (65 mg/kg i.p.) and the rats’ hind paw reaction was examined fifteen minutes later to see if they were completely anesthetized(Gage et al., 2012). Followed by transcranial perfusion with ice-cold 0.9% NaCl, decapitated, and the brain was removed.

### 2.8 IHC staining

Briefly, paraffin slices were deparaffinized in Xylene and rehydrated using graded alcohols, as previously described(Forsyth et al., 2011). They were then subjected to microwave for 10 min with citrate buffer followed by cooling to room temperature. Slides were washed with 0.1% PBS and blocked for 30 min with Normal Goat Serum (Maixin, Shanghai). The cells were then treated with primary antibodies overnight at 4 °C and subsequently incubated with secondary antibody for an hour at RT and permeabilized in xylene and mounted. All of the photographs were taken with a Zeiss AXIO Imager. Z1 and a digital AxioCam HRm, and then processed with AxioVision 4.8 and Adobe Photoshop

### 2.9 HE staining

The HE staining was carried out according to standard procedure(Ren et al., 2007). Sections (5 μm) were stained with hematoxylin solution for 5 min, 1% acid ethanol for 15 s, washed in distilled water, stained with eosin for 7 min, dehydrated with graded alcohol, and cleared in xylene after being deparaffinized and rehydrated. For microscopic inspection, the slides were evaluated in Vectra3 (Akoya BIOSCIENCES, Shanghai).

### 2.10 Western Blot analysis

The Western Blot analysis was carried out as stated previously(Hirano, 2012; MacPhee, 2010). In a nutshell, RIPA buffer was used to obtain total protein extracts. SDS-PAGE was used to evaluate equal amounts of proteins, which were then transferred to polyvinylidene difluoride membranes. The membranes were blocked for 2 h at room temperature with a solution of 5% nonfat dry milk in TBST. The membranes were then treated with particular primary antibodies overnight at 4 °C. The membranes were treated with secondary antibodies for 2 h at room temperature the next day. The western blot analysis was performed using the ECL chemiluminescence method.

## 3 Statistical analysis

SPSS 16.0 Software (SPSS, Chicago, IL, USA) and GraphPad Prism 6.0 software (GraphPad Software Inc. La Jolla, CA) were used to evaluate the results which are expressed as mean ± SD.

A one-way analysis of variance (ANOVA) was employed, followed by Tukey’s HSD post hoc test and Holm-Bonferroni correction for multiple comparisons between groups. Two-way repeated-measures ANOVA was used to analyze the neurobehavioral functions over time. P<0.05 was considered statistically significant.

## 4 Result

### 4.1 Ifenprodil can improve differently the effects on animal behavior

Time points and treatment regimen are shown in Fig.1. We selected 6-OHDA-PD rats and studied the effects of systemic administration (intraperitoneal injection, i.p.) versus brain injection. We also investigated the striatum and the substantia nigra, two brain regions that are tightly linked to dopamine and glutamatergic systems.

**Fig 1.**
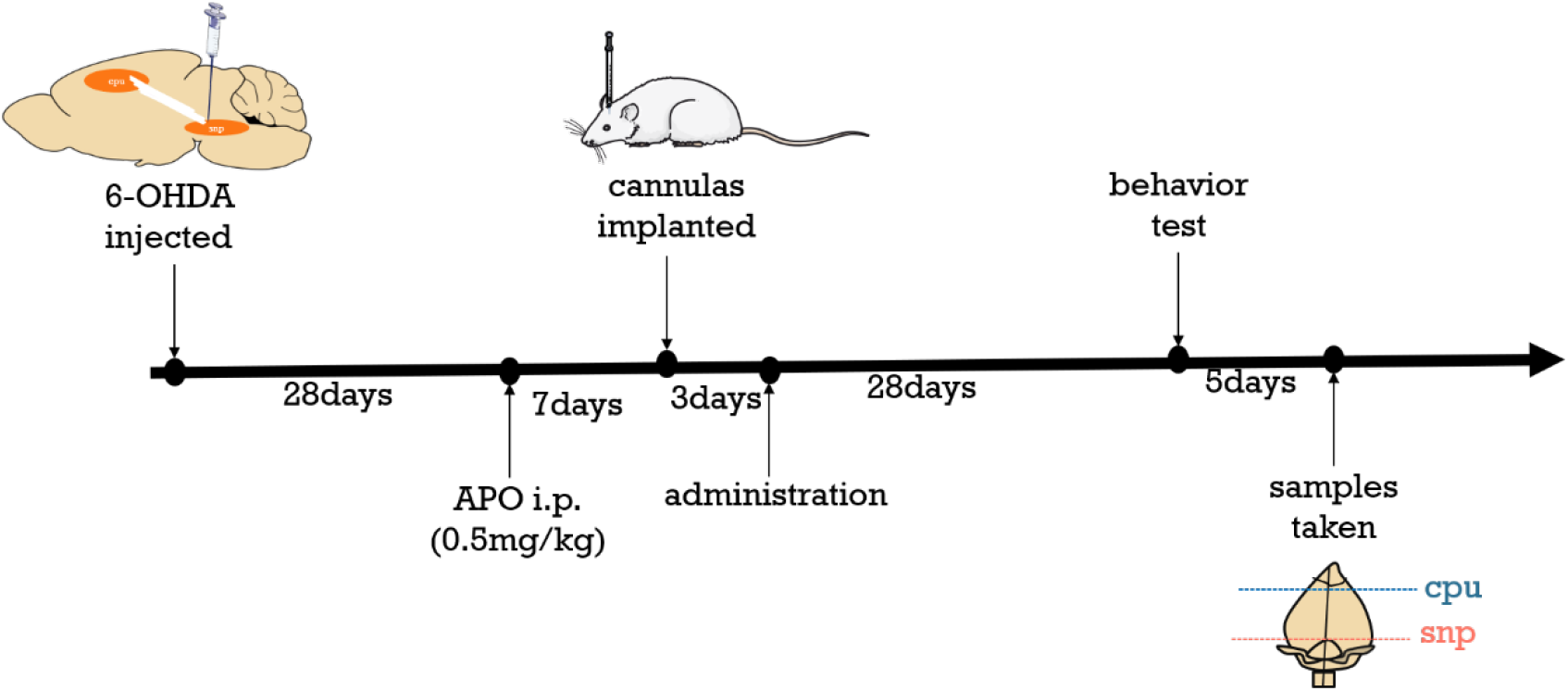
Flow chart of animal experiments (n = 12, each group). cpu: abbreviation for caudate putamen (striatum); snp: abbreviation for substantia nigra part; i.p. : abbreviation for intraperitoneal administration.

### 4.2 Rotarod test

All rats were placed on a rotarod apparatus that accelerated from 0 to 30 rpm to determine the retention time on the revolving rod after different treatments to test their motor coordination. As shown in Fig.2, over time, Ifenprodil continuously enhanced 6-OHDA-lesioned rats of the time staying on the rod in all drug delivery. Compared with the model, CPU begins to diverge on 14^th^ and shows a significant difference (p < 0.05) and SNR shows a significant difference on 14^th^ and 17^th^ (p < 0.05). Upon comparison with i.p. group, CPU shows significant difference on 14^th^ and 28^th^ (p < 0.05) and SNR didn’t show any difference significance in the whole data set. We also analyzed two different brain injection regions, we found that CPU and SNR only show a different significance on 14^th^ (p< 0.05), therefore we cannot individually evaluate the effects of the two brain injection regions.

**Fig 2.**
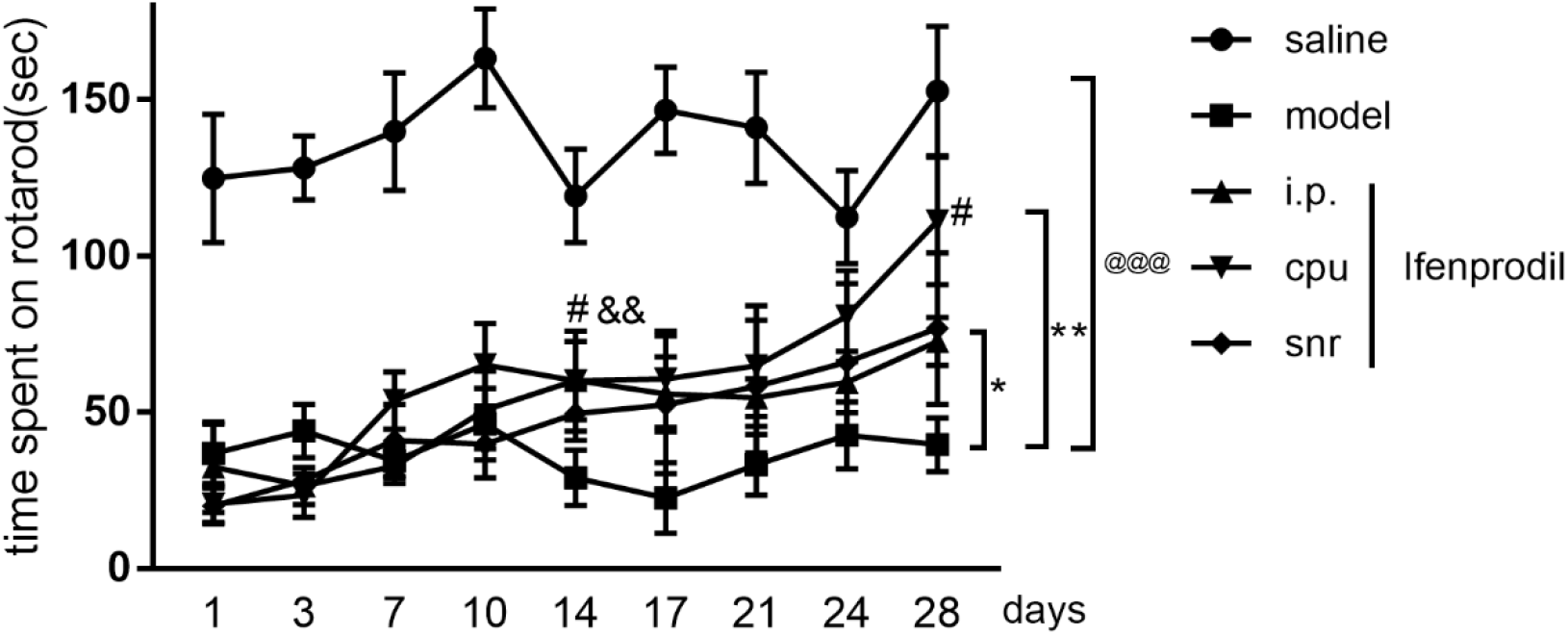
The latency to fall off from a rotating drum in experimental rat (n = 12). Mean ± SEM is used to represent the data. ^@^p < 0.05, ^@@^p < 0.01 and ^@@@^p < 0.001 versus saline group, ^*^p < 0.05, ^**^p < 0.01 and ^***^p < 0.001 versus saline group, ^#^p < 0.05 versus i.p. group, ^&^p < 0.05 versus CPU group. Student’s t-test.

### 4.3 Open field

The open-field experiment was used as an important way to evaluate athletic ability. A high behavioral frequency indicates increased locomotion and exploration. The results of the open-field test are shown in Fig. 3, where animals in the Ifenprodil-treated groups had a higher frequency of behavior than the model, and the CPU indicated a significant difference in contract relative to the model (p<0.05). Although we did not see significant differences between i.p. or SNR with the model, we cannot deny the therapeutic effect of other treatments because they showed the therapeutic tendency.

**Fig 3.**
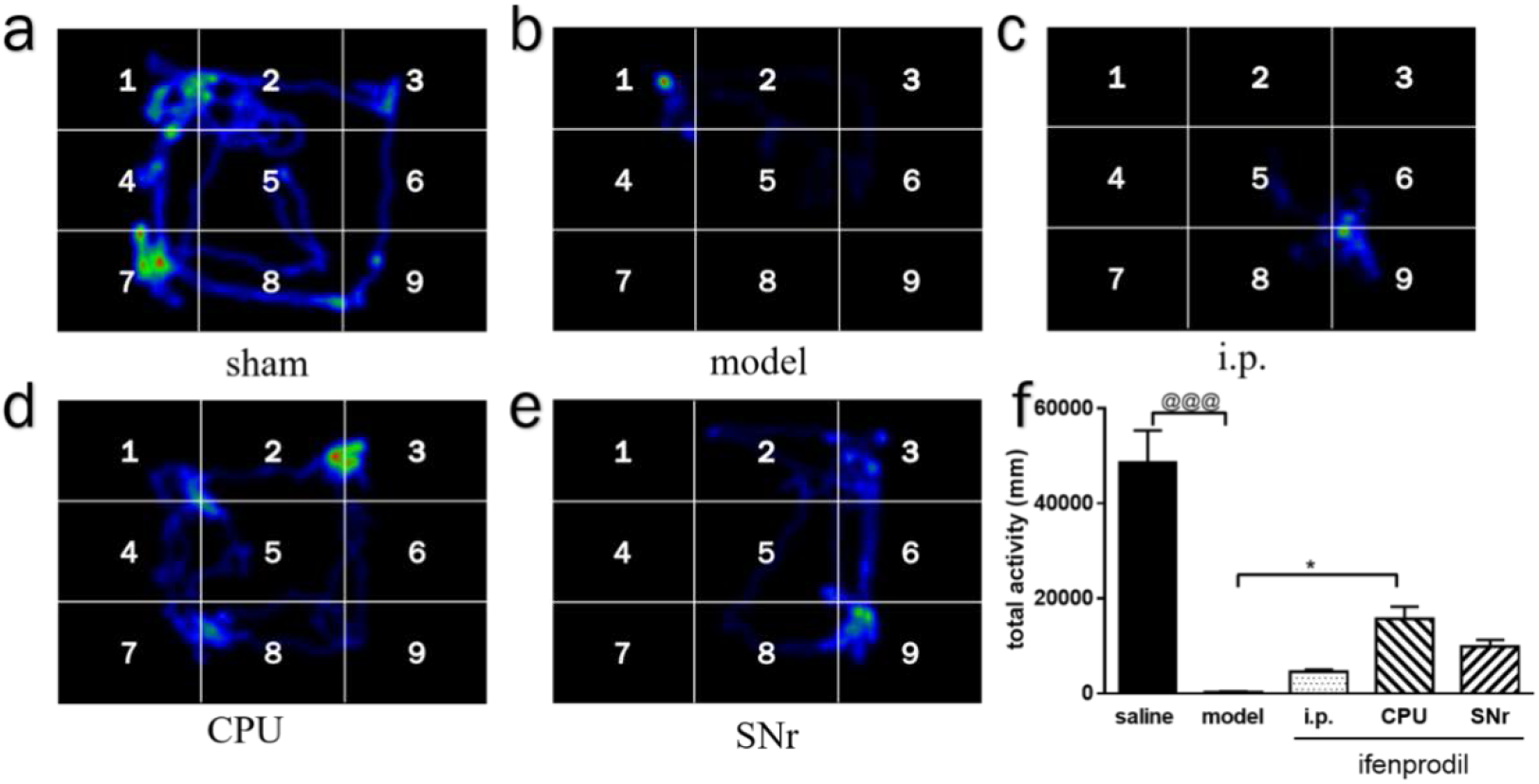
Open-field behavior. (a~e). Heat maps of rats exploring behavior in the open field test (n = 12). (f). Locomotor activity in open-field test over 10 min in experimental rat (n = 8). Mean ± SEM is used to represent the data. ^@^p < 0.05, ^@@^p < 0.01 and ^@@@^p < 0.001 versus saline group, ^*^p < 0.05, ^**^p < 0.01 and ^***^p < 0.001 versus model group, ^#^p < 0.05 versus i.p. group, ^&^p < 0.05 versus CPU group. Student’s t-test.

### 4.4 Ifenprodil treatment shows a protective effect on neurons both in substantia nigra and striatum

Brains were taken, fixed in 10% buffered formalin, chopped into slices, immunostaining was conducted to identify dopaminergic (tyrosine hydroxylase, TH) and sate of microglia, and HE staining revealed neuronal morphologies in the Substantia nigra and Striatum of rats after the last behavior test.

The results of HE staining are shown in Fig. 4, in control section of the substantia nigra compact part, the density of neurons is equally distributed and dense, color is clearly identifiable, neurons are intact and are large/medium-sized pyramidal cells, the arrangement is regular, and there is no degeneration or necrosis. 6-OHDA-lesioned neurons were smaller, had an incomplete shape, were ambiguous in outline, and had a much lower number of neurons. The number of necrotic cells in the striatum was reduced after 3 weeks of therapy, with CPU treatment having the best effect (p<0.05), morphology and structure returning to normal, and the same outcomes were seen in the substantia nigra.

**Fig.4.**
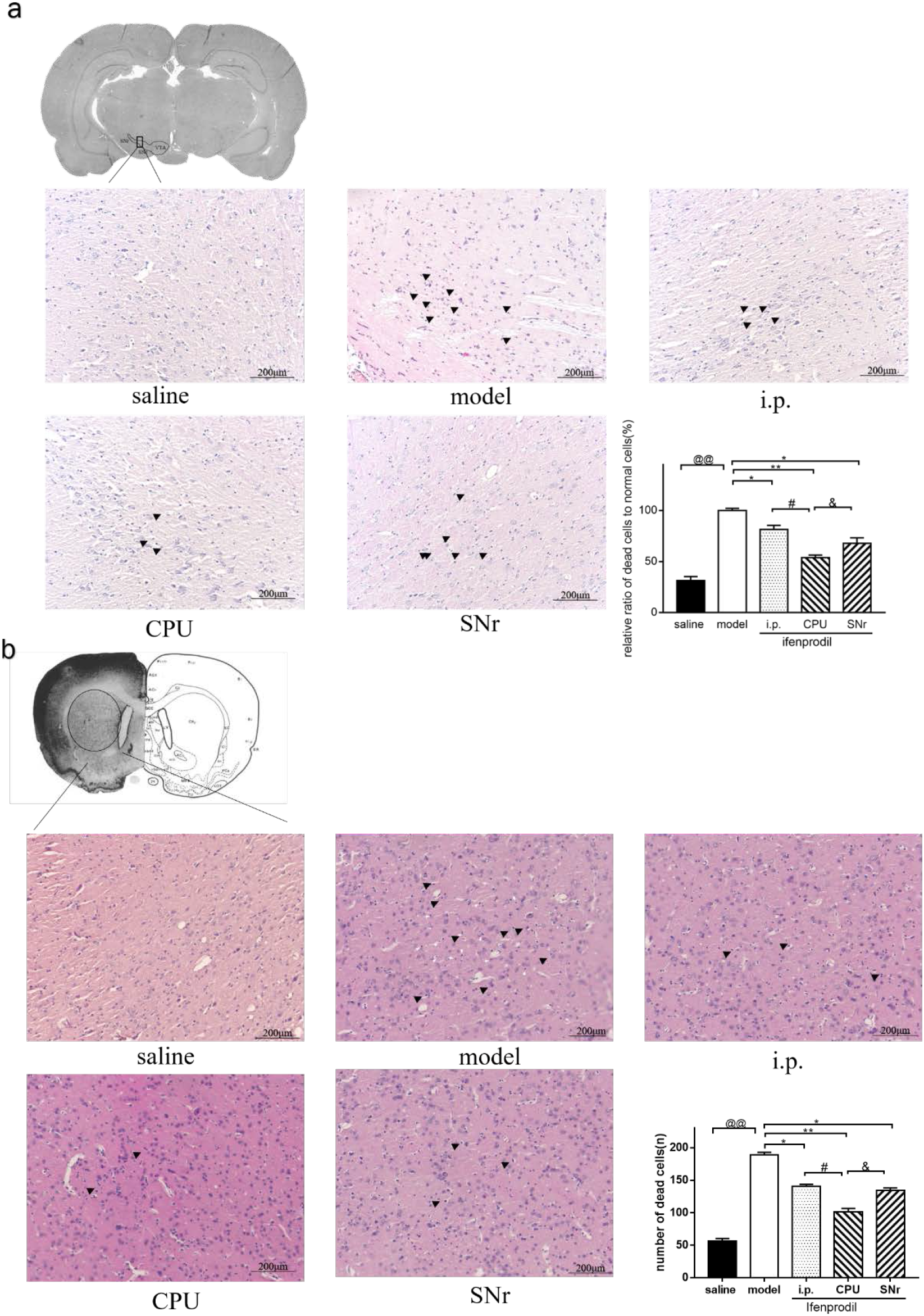
HE staining. (a). Substantia nigra (n=8). (b). Striatum (n = 8). (c). The number of apoptotic cells in the striatum. Black arrows point to necrotic pyramidal cells. Randomly selected visual fields were counted in each group. Mean ± SEM is used to represent the data. ^@^p < 0.05, ^@@^p < 0.01 and ^@@@^p < 0.001 versus saline group, ^*^p<0.05, ^**^p<0.01 and ^***^p<0.001 versus model group, ^#^p<0.05 versus i.p. group, ^&^p<0.05 versus CPU group. Student’s t-test.

In Fig.5 (b), we compared microglia activation in the striatum, and the results revealed that Ifenprodil reduces microglia activation to relieve inflammation, and CPU seems to have a significant effect (p<0.05). Simultaneously, we evaluated the functional viability of dopaminergic neurons in the striatum, as well as the expression of the rate-limiting tyrosine enzyme using an anti-TH antibody (Fig. 5) (a). Ifenprodil protected TH neurons, reduced apoptosis in neurons, with a significant effect of CPU (p<0.05)

**Fig.5.**
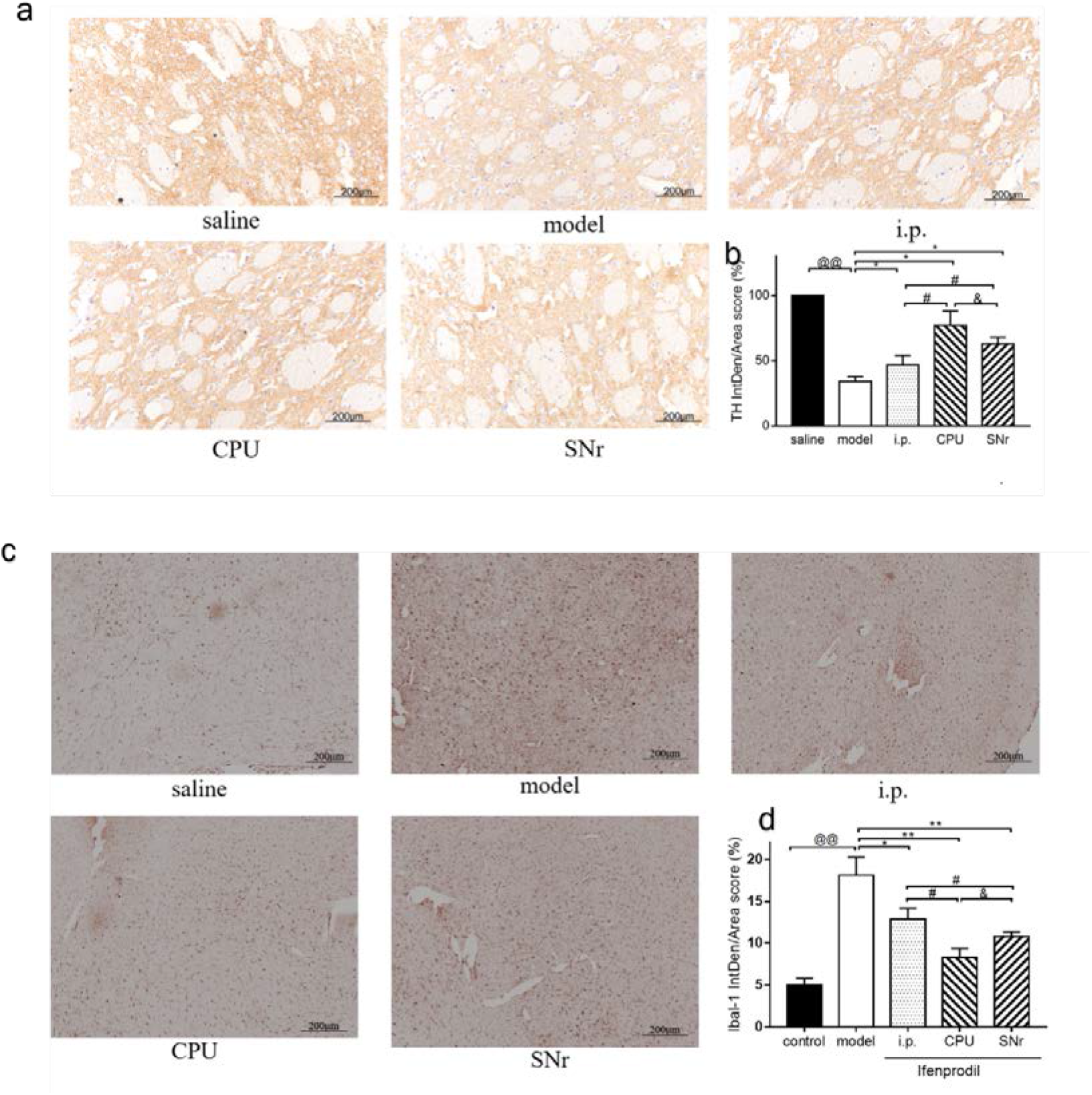
IHC staining. (a-b) Optical density of TH-positive cells in the striatum (n = 8). (c-d) Optical density of ibal-2 in the striatum (n = 8). Mean ± SEM is used to represent the data. ^@^p < 0.05, ^@@^p < 0.01 and ^@@@^p < 0.001 versus saline group, ^**^p<0.01 and ^***^p<0.001 versus model group, ^#^p<0.05 versus i.p. group, ^&^p<0.05 versus CPU group. Student’s t-test.

### 4.5 Ifenprodil changed NMDA receptor subunits expression in the striatum

To see if 6-OHDA-lesioned and Ifenprodil influence the expression of NMDA receptor subunits, and to see if PSD95/NMDAR2B-CaMKII pathways are involved using western blot analysis. We assessed, in Fig.6, the levels of postsynaptic membrane NMDAR2B, pho-NMDAR2B (1303), and pho-NMDAR2B (1472) in the striatum, and the changes in proteins in the PSD95/NMDAR2B-CaMKIIsignaling pathway. The results revealed that there was no difference in total NMDAR2B protein concentration between the groups, but S1303 phosphorylated and T1472 phosphorylated indicated an increase in model size. We didn’t see any significant differences in the expression level of CaMKII and α-Ca protein in each group, but CaMKII-phosphorylated and β-Ca showed an increase in the model. Ifenprodil had a good effect in every treated group and CPU showed a significant decrease in PSD95, S1303 phosphorylated, T1472 phosphorylated, phosphorylated-CaMKII, and β-Ca.

**Fig.6.**
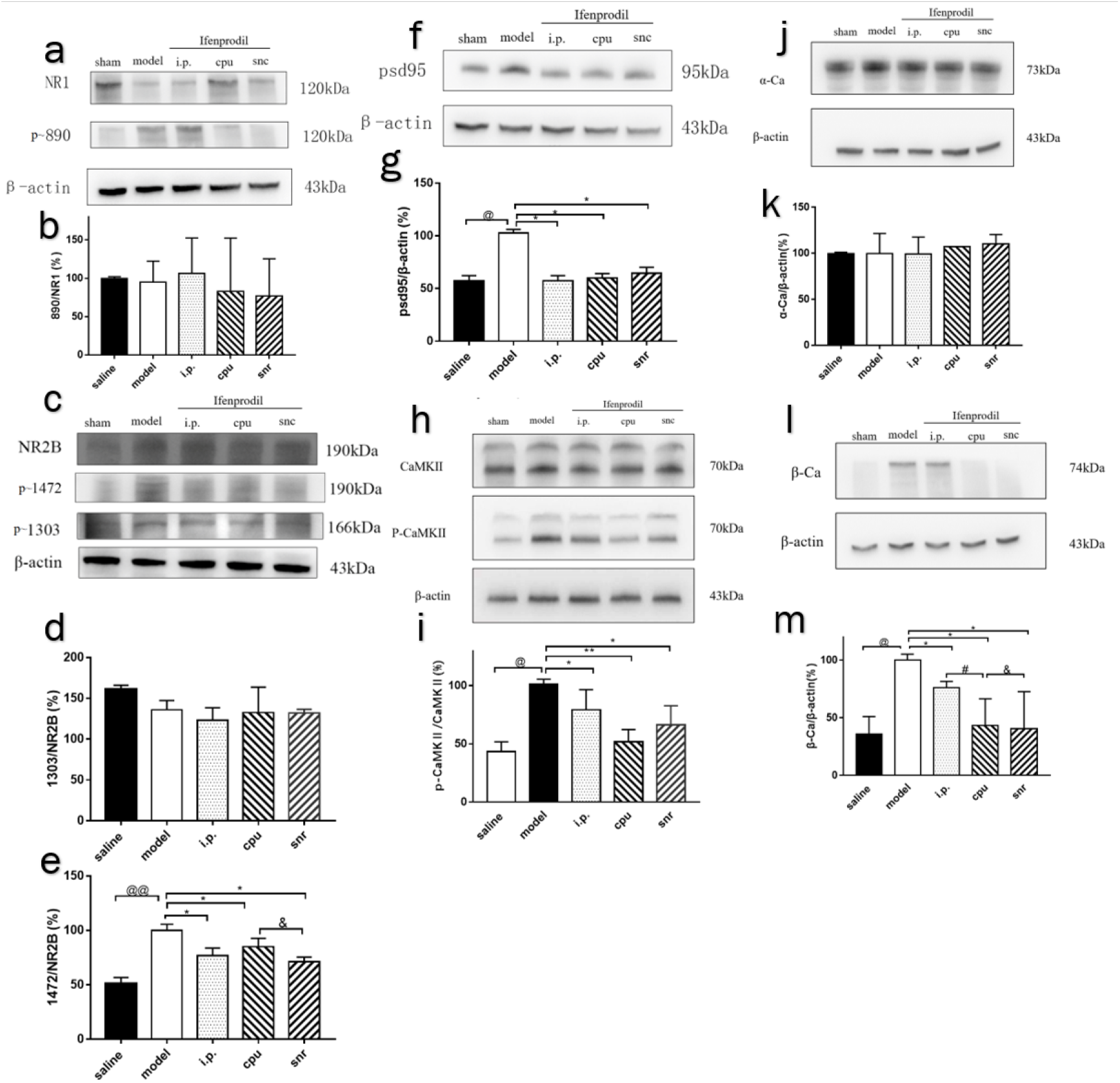
Protein-expression measurement. (a-e) Western blot and densitometry analysis of NMDA receptor (n = 8). (f-g) A quantitative analysis of the relative expression levels of psd95 (n = 8). (h-m) The densitometry of calcium signal-related protein expression in the midbrain compared to that of β-actin is displayed in striatum tissue (n = 8). Mean ± SEM is used to represent the data. ^@^p < 0.05, ^@@^p < 0.01 and ^@@@^p < 0.001 versus saline group, ^*^p<0.05, ^**^p<0.01 and ^***^p<0.001 versus model group, ^#^p<0.05 versus i.p. group, ^&^p<0.05 versus CPU group. Student’s t-test.

### 4.6 Involvement of the autophagy pathway in the neuroprotective effect of Ifenprodil

To assess autophagy activity, we looked at changes in autophagic and apoptotic proteins in the striatum and midbrain at the same time in Fig.7. The results showed that 6-OHDA-lesioned striatum induced a significant drop in lc3b, beclin-1, and bcl-2, as well as an increase in p62, bax, and PARP, whereas treatments tended to raise lc3b, beclin-1, and bcl-2, while decreasing p62, bax, and PARP. BNIP3L/Nix, a mitochondrial autophagy receptor that promotes mitochondrial autophagy by binding to LC3 during cell development or pathological circumstances, also showed an increase in treatment tendencies. However, autophagy inhibition in the midbrain of 6-OHDA-lesioned rats is demonstrated by a decrease in lc3b, beclin-1, and bcl-2, as well as an increase in p62, following the end of Ifenprodil treatment.

**Fig.7.**
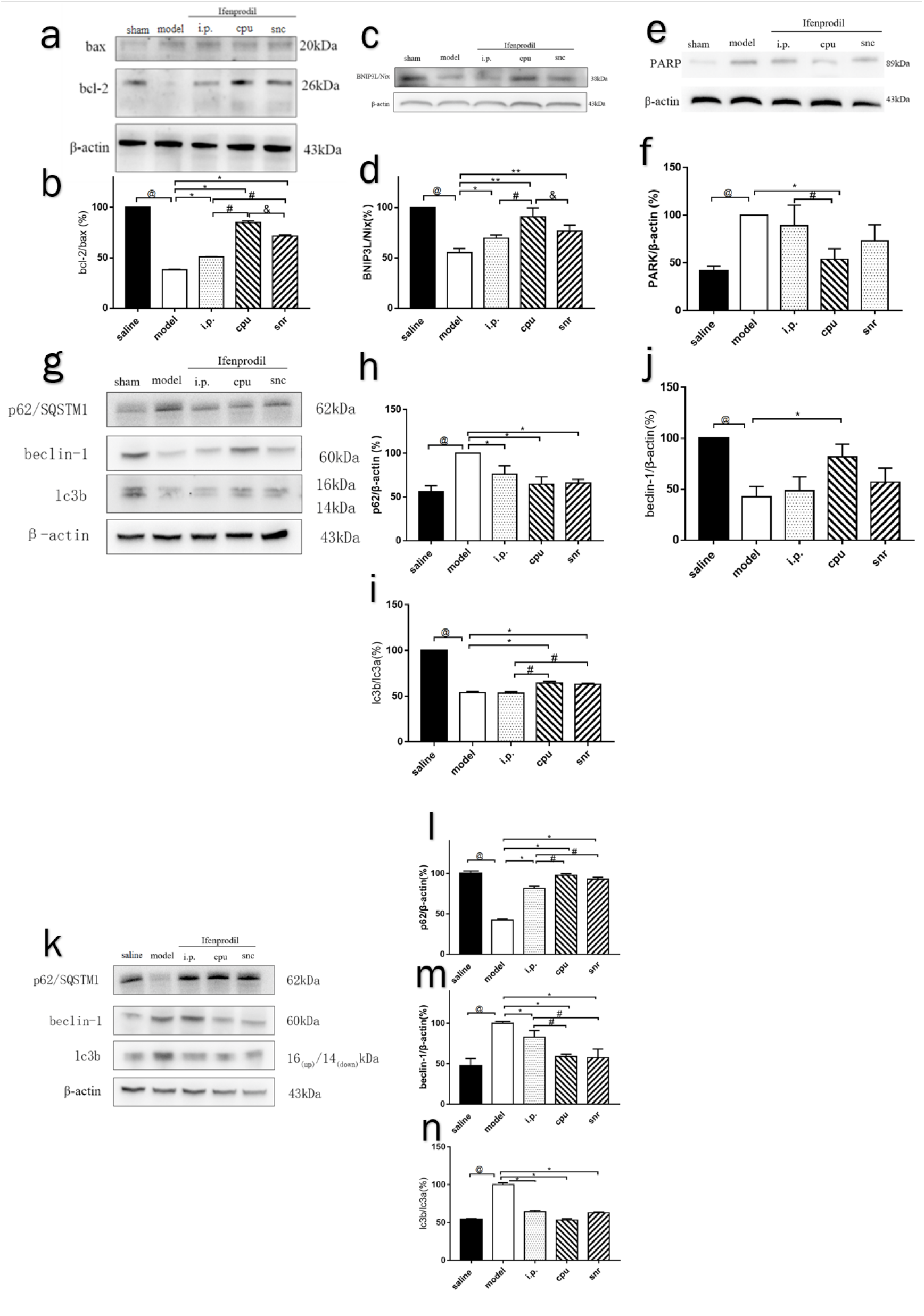
Protein expression. (a-j) Western blot and densitometry analysis of striatum tissue (n = 8). (k-n). The densitometry of autophagy-related protein expression in the midbrain compared to that of β-actin is displayed of midbrain tissue (n = 8). Mean ± SEM is used to represent the data. ^@^p < 0.05, ^@@^p < 0.01 and ^@@@^p < 0.001 versus saline group, ^*^p<0.05, ^**^p<0.01 and ^***^p<0.001 versus moel group, ^#^p<0.05 versus i.p. group, ^&^p<0.05 versus CPU group. Student’s t-test.

## 5 Discussion

As it shows in Fig.8, the large rise in intracellular Ca^2+^ elicited by glutamate-induced NMDA receptor activation serves a key role in initiating intracellular events that lead to cell death. NR2B antagonists have previously been shown to have therapeutic effects in PD animal models(Gogas, 2006b; Nutt et al., 2008). To examine the mechanism of NR2B involvement under 6-OHDA-lesioned, we used Ifenprodil as a protective drug in this experiment. For instance, Ifenprodil has a ~400-fold greater affinity for the 2B subunit than the 2A subunit(Gogas, 2006a). Then, as a PD animal model, we chose 6-OHDA rats to investigate the link between NR2B and glutamate excitotoxicity.

**Fig.8.**
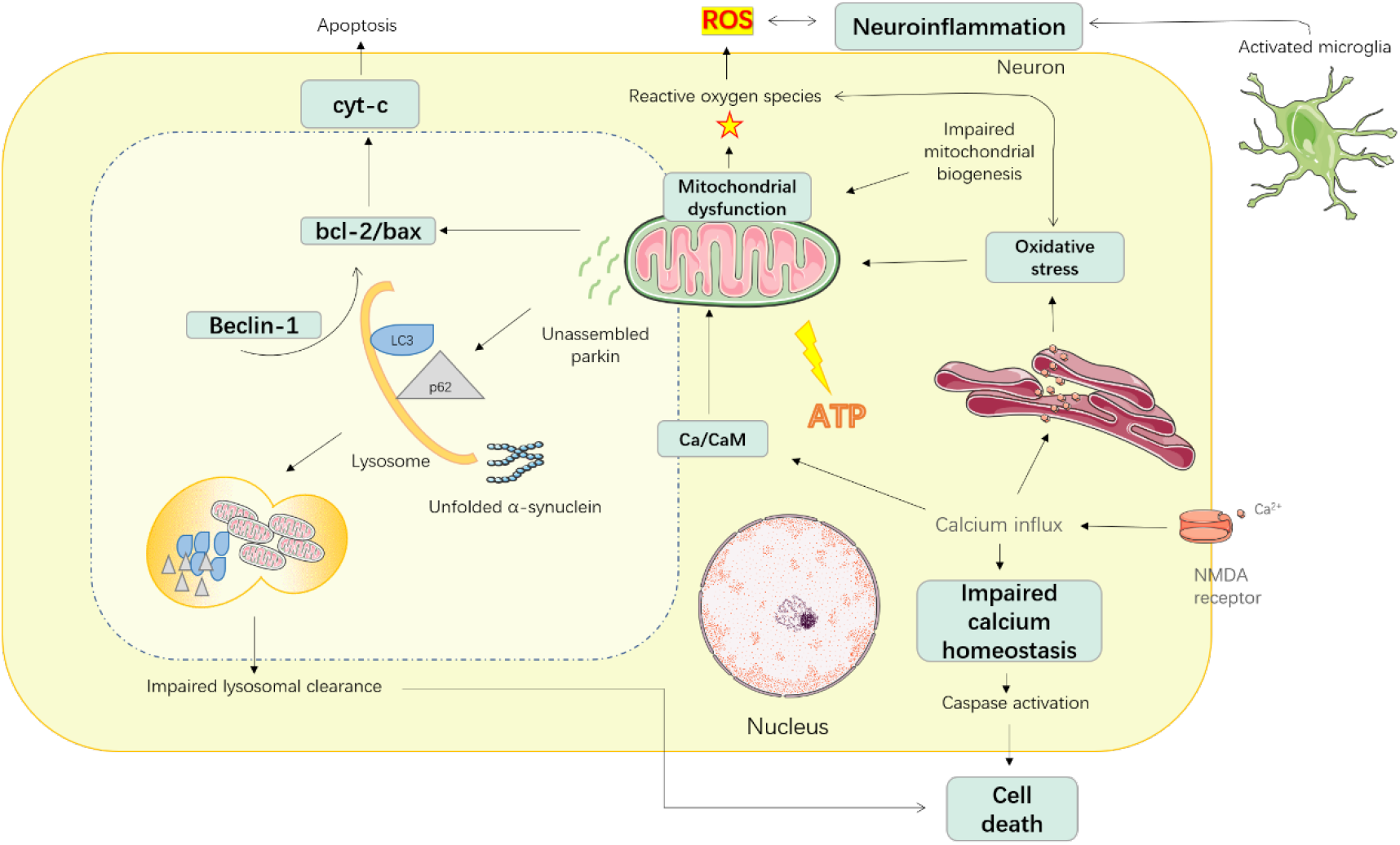
Diagram of the mechanism of overload-released glutamate triggering neuronal death by overactivation of NMDA receptors in PD. Glutamate, the major excitatory neurotransmitter in the central nervous system, is known to induce excitotoxicity by binding to the NMDA receptor, then triggers a calcium influx that activates calpain and induces mitochondrial dysfunction. Damaged mitochondria lead to ATP depletion, ROS production, and further disruption of calcium homeostasis which is implicated in neuronal death in PD. Autophagy can clear damaged and senescent mitochondria, but the recognition of autophagosomes to the damaged mitochondria is interrupted in PD, and the impaired mitochondria cannot be delivered to lysosomes by autophagosomes causing the injured mitochondria in cells cannot be cleared in time. Neuronal death.

We used an in vivo 6-OHDA-induced PD model in rats. We know that glutamatergic neurons can project to the anterior and posterior striatum, as well as the substantia nigra, from various areas of the prefrontal cortex(Blandini et al., 1996). An increase in glutamate receptor density in the rat striatum after a 6-OHDA lesion or in PD patients would support the idea of increased glutamatergic synaptic activity(Chotibut et al., 2014). A decrease in synaptosomal Glu uptake was reported in 3 weeks post-6-OHDA-lesioned rats, that a significant increase in the Glu levels were found in the striatum (156% higher than normal)(Iovino et al., 2020; Wang et al., 2020). In this study, we used systemic administration and stereotactic injection into the brain to assess Ifenprodil’s ability to protect against the neurotoxic effects of 6-OHDA. As previously reported, the output signal of dopamine cells in the midbrain (particularly the SNr) to the striatum (particularly the dorsal striatum) is related to movement; that is, dopamine signals increase before the mouse begins to exercise from rest and after the acceleration during exercise; thus, intracerebral administration was used to inject into the striatum and substantia nigra pars reticulata(Lindgren et al., 2010; Weise et al., 2009).

We tested motor coordination by rotarod test and activities by an open field, results showed that Ifenprodil can improve the exercise capacity of rats, and the effect of brain administration was better than systemic administration, CPU group showed the best improvement among all. We got almost the same treatment conclusion in the HE results. To investigate the neuroprotective effects of Ifenprodil, we measured the expression of ibal-2 and TH positive neurons, as well as the expression of autophagy protein and NMDA receptors in these samples. Among them, ibal-2 can be used to detect both resting and activated microglia, and activated microglia is a cellular hallmark of neuroinflammation(Kang et al., 2019). TH is a dopaminergic neuron marker that can, to some extent, indicate the number of DA neurons. Based on the findings, we can conclude that Ifenprodil could reduce microglial activation while also protecting TH-positive neurons.

Then we looked into the autophagy issue in relation to PD. LC3 is a significant autophagy marker among them. When autophagy is established, cytoplasmic LC3 will enzymatically digest tiny peptides and convert to membrane type (ie LC3-b). The LC3-b/a ratio can be used to assess autophagy. Beclin-1 and p62 are involved in autophagy; when autophagy is suppressed, inactivated beclin-1 causes a decline in the ability of cells to regulate autophagy, which aggravates the apoptotic process; in the meantime, p62 undergo accumulation(Mishra et al., 2018; Schmitz et al., 2016). We analyzed the expression of autophagy proteins in the striatum and midbrain. The results showed that Ifenprodil administration could increase the expression of LC3b, beclin-1, BNIP3LNix, and bcl-2, while inhibiting the expression of p62, bax, PARP in the striatum, which suggested that Ifenprodil improved the inhibition of the autophagy pathway in the striatum, interestingly, the results in midbrain showed opposite, that Ifenprodil inhibited the enhancement in the midbrain. Furthermore, we discovered that NR2B expression in the striatum of PD rats is extremely high, but there is no expression in the substantia nigra region, which is consistent with previous reports(Jin et al., 1997; Wang et al., 1995).

Dopamine neurons in the brain are found in the substantia nigra and project to the striatum. The striatum is a huge enrichment area of NMDA receptors and is the input structure of the basal ganglia, which is located below the forebrain cortex(Jin et al., 1997). In general, the degradation of dopaminergic neurons in the substantia nigra compact area of PD patients results in an overabundance of STN glutamatergic neurons projecting to the basal ganglia’s input structure, as well as a large accumulation of glutamine in the striatum, resulting in the post-synaptic membrane. Excessive excitation activates NMDA receptors, allowing a significant amount of Ca^2+^ to enter the cell. The mitochondrial-autophagy pathway is inhibited by severe calcium overload, which damages mitochondrial activity. However, neuronal loss in 6-OHDA-damaged dopamine neurons in the substantia nigra was not linked to NMDA receptors, which could be due to the direct influence of toxins.

In conclusion, we show that NR2B is implicated in the course of PD and that Ifenprodil has a neuroprotective impact. The CaMKII-Autophagy pathway is influenced by NR2B, and the NMDA receptor’s activation role in PD is linked to autophagy, which is mediated by calcium ions entering the NMDA receptor-associated channel. As a result, NMDA receptor antagonists like Ifenprodil could be useful in the treatment of PD.

## 6 Funding

This project was supported by Yantai University. The author(s) disclosed receipt of the following financial support for the research, authorship, and/or publication of this article: This work was supported by the “Taishan Industry Leading Talent Laureate”.

## 7 Acknowledgements

Xinyu ZHAO performed the research, analyzed the data, and wrote the manuscript. Fufang TIAN contributed to animal experiments. Xin YU designed and funded the research, interpreted the data, finally approved the submission of this manuscript and revised the manuscript.

## Notes

### Competing Interest Statement

The authors have declared no competing interest.

